# The Olivary Pretectal Nucleus Receives Visual Input of High Spatial Resolution

**DOI:** 10.1101/2020.06.23.168054

**Authors:** Jared N. Levine, Gregory W. Schwartz

## Abstract

In the mouse, retinal output is computed by over 40 distinct types of retinal ganglion cells (RGCs) (Baden et al., 2016). Determining which of these many RGC types project to a retinorecipient region is a key step in elucidating the role that region plays in visually-mediated behaviors. Combining retrograde viral tracing and single-cell electrophysiology, we identify the RGC types which project to the olivary pretectal nucleus (OPN), a major visual structure. We find that retinal input to the OPN consists of a variety of intrinsically-photosensitive and conventional RGC types, the latter a diverse set of mostly ON RGCs. Surprisingly, while the OPN is most associated with the pupillary light reflex (PLR) pathway, requiring information about absolute luminance, we show that the majority of the retinal input to the OPN is from single cell type which transmits information unrelated to luminance. This ON-transient RGC accounts for two-thirds of the input to the OPN, and responds to small objects across a wide range of speeds. This finding suggests a role for the OPN in visually-mediated behaviors beyond the PLR.

**Significance statement:** The olivary pretectal nucleus is a midbrain structure which receives direct input from retinal ganglion cells (RGC), and modulates pupil diameter in response to changing absolute light level. In the present study, we combine viral tracing and electrophysiology to identify the RGC types which project to the OPN. Surprisingly, the majority of its input comes from a single type which does not encode absolute luminance, but instead responds to small objects across a wide range of speeds. These findings are consistent with a role for the OPN apart from pupil control and suggest future experiments to elucidate its full role in visually-mediated behavior.

## Introduction

The olivary pretectal nucleus (OPN), one of several retinorecipient nuclei of the pretectum, has been studied for decades as a key structure in the pupillary light reflex (PLR) pathway, given its ability to encode luminance information (Trejo and Cicerone, 1984; Clarke and Ikeda, 1985) and its connectivity with the downstream effectors of the pupil reflex (Klooster *et al.*, 1995; (Klooster et al., 1993). Previous findings have indicated that the retinal projection to the OPN is bilateral and substantial (Young and Lund, 1998; Morin and Studholme, 2014), but while some of the RGC types projecting to the OPN have been identified, there has been no comprehensive classification of OPN-projecting RGCs. Characterization of transgenic mouse lines has identified projections to OPN from subsets of RGCs, but these findings are complicated by the fact that multiple RGC types are labeled in these lines (Rousso et al., 2016; Martersteck et al., 2017). A number of studies have described a strong projection of intrinsically-photosensitive (ip) ganglion cells to the OPN, suggesting that this class of RGC forms the predominant portion of total retinal input to this structure (Hattar et al., 2006). More recent work has identified selective innervation of the OPN shell region by a subset of M1 ipRGCs as critical for a fully functional PLR (Chen, Badea and Hattar, 2011). However, this work also indicates a non-melanopsin-expressing set of RGCs projects to the OPN core. Furthermore, anatomical studies have described connectivity between the OPN and brain regions that are involved in vision but not part of the canonical PLR circuit, such as the superior colliculus, the nucleus of the optic tract, and the ventral lateral geniculate nucleus (Klooster *et al.*, 1995). Thus, the OPN – particularly its core region – may play a role in visual behavior beyond the PLR, and identifying the complement of RGCs projecting to this structure represents an important step in uncovering the multiplexed and parallel code from the retina to the brain.

We combined retrograde viral tracing with single-cell electrophysiology to identify the RGC types which project to the OPN. We found projections from a large variety of RGC types, both ipRGCs and conventional RGCs. The majority of input to the OPN was carried by a single RGC type, and while it has been molecularly identified as an ipRGC called the M6 (Quattrochi et al., 2019), we found that it does not encode absolute luminance. Rather, it is a transient ON RGC that responds to small stimuli across a wide range of speeds. These results suggest a special role for M6 RGCs in transmitting information to OPN that could be used in the spatiotemporal processing of small moving objects, augmenting our understanding of the visual function of this brain region.

## Methods

### Animals

Wild-type mice of both sexes of age P60-90 were used for all experiments.

### Stereotactic virus injections

Prior to surgery, anesthesia was induced with the inhalable anesthetic isoflurane (∼3% diluted in oxygen). During the procedure, anesthesia was maintained with ∼1.5% isoflurane. Meloxicam was administered subcutaneously to reduce pain and edema (1.5 mg/kg in 10% saline). A small circular craniotomy was centered over the OPN using the following coordinates to target the rostral aspect: −2.50mm anterior/posterior, 0.60 mm medial/lateral and −2.37 mm dorsal/ventral. Subsequently, AAV engineered for retrograde transport and carrying either tdTomato or green fluorescent protein (AddGene, AAVrg-CAG-tdTomato, AAVrg-CAG-GFP) in PBS was injected using a Nanoject II (Drummond) fitted with a glass pipette with an inner diameter of 10–20 μm. 4 pulses of 9.2 nL each (36 nL total volume), at 30 s intervals, were delivered. Following injection, mice were kept alive for 5-7 days to allow for sufficient fluorescent protein expression in retinal ganglion cell bodies. The injection site was confirmed in post hoc in brain slices according to anatomical landmarks, or with intravitreal injection of 2μl of the cholera toxin b subunit (Sigma-Aldrich) into the contralateral eye.

### Retinal dissection and preparation

Wild-type mice of either sex were dark-adapted overnight. Dissection and excision of retinal tissue were performed under infrared illumination (900 nm) using night-vision goggles and night-vision dissecting scope monocular attachments. Research animals were sacrificed in accordance with all animal care standards provided by Northwestern University. The retina was placed onto a glass coverslip coated with poly-d-lysine, with the ganglion cell layer facing upward. The retina was superfused with carbogenated Ames solution warmed to 32°C (US Biological Life Sciences, A-1372-25).

### Immunohistochemistry and histology

Retinas were fixed for 10 min in 4% paraformaldehyde (Electron Microscopy Sciences) and incubated in 0.1 M phosphate buffer (PB) overnight at 4 °C. Fixed retinas containing cells filled with neurobiotin were then incubated in PBS containing 3% normal donkey serum (blocking agent), 0.05% sodium azide, 0.5% Triton X-100 overnight. This was followed by incubation in blocking solution and primary antibody against ChAT (Millipore, AB144P, goat anti-ChAT, 1:500 v/v) for five nights at 4 °C. Afterward, tissues were rinsed in 0.1 M PB and incubated for two nights at 4 °C with a secondary antibody against goat IgG (Jackson ImmunoResearch, 711-605-152, donkey anti-goat Alexa 647, 1:500 v/v) and streptavidin (Thermo Scientific, DyLight 488, 1:500 v/v). Following immunostaining, retinas were mounted on slides with Vectashield Antifade mounting (Vector Labs) medium. Following extraction, brains were fixed overnight in 4% paraformaldehyde (Electron Microscopy Sciences). Fixed brains were then frozen in O.C.T. (Sakura, 4583) before sectioning. Frozen sections of 30 μm were cut for identification of the injection site.

### Imaging

Fixed tissues were imaged on a Nikon A1R laser scanning confocal microscope mounted on a Nikon Ti ZDrive PerfectFocus microscope stand equipped with an inverted ×40 oil immersion objective (Nikon Plan Apo VC ×40/×60/1.4 NA). RGC dendrites and ChAT labeling were imaged at 488 and 647 nm excitation, respectively. Dendritic arbors were traced using Fiji (ImageJ) software with the Simple Neurite Tracer plugin. Dendrites were traced and computationally flattened relative to the ChAT bands (Sümbül et al., 2014). Epifluorescence images of brain sections and retinas were acquired on a Nikon Ti2 widefield microscope.

### Visual stimuli

Visual stimuli were presented with a custom-designed light-projection device capable of controlling patterned visual stimulation at high frame rates (1.4 kHz). Stimuli were generated on a 912 × 1140-pixel digital projector using the blue (450 nm) LED and focused on the photoreceptors. We report light intensities in rod isomerizations (R*) per rod per second. Based on the spectrum of our blue LED, the spectral sensitivities of mouse opsins, and collecting areas of mouse photoreceptors, each R* corresponds to 0.3 isomerizations per M-cone opsin and 6 × 10^−4^ isomerizations per S-cone opsin. Stimuli were first aligned to the RF center of each cell using a series of flashing horizontal and vertical bars from darkness (200 × 40 μm) across 13 locations along each axis, spaced 40 μm apart. Spots of diameters ranging from 10–1200 μm from darkness were used to characterize the spatial dynamics of RGC responses. For M6 RGC receptive field mapping, we used a spot of 40 μm diameter flashed from darkness at different locations. Moving bars were presented from darkness at three orientations, and were 1000 μm long, and varied in width and drifting speed. Stimuli from darkness were presented at 200 rod isomerizations (R*)/rod/s.

### Analysis

Analysis of electrophysiology data was done with custom Matlab code (https://github.com/SchwartzNU/SymphonyAnalysis). Peak firing rates of M6 RGCs to the moving bar stimulus were calculated from peri-stimulus time histograms of spiking responses. For calculation of M6 receptive field diameter, first a spatial map of spike rate vs stimulus position was generated. Responses to individual spots were separated and peak values were averaged to generate a value for each position. These values were displayed on the grid locations to create a 2D RF strength map. A two-dimensional Gaussian fit was applied, the standard deviation in x and y were treated as the axes of an ellipse, and the area of the ellipse calculated as *π*\**a*\**b*. This area was then treated as circular, and the diameter calculated accordingly. For RGC soma counting to determine peak density of labeled RGCs, a square 1 mm x 1 mm was drawn in the region of densest labeling, as determined by eye. The number of labeled cells was then determined using the ImageJ Objects Counter plugin.

### Statistical analysis

Statistical analysis was performed using Igor Pro 8 (WaveMetrics) and MATLAB (MathWorks). Plots show means and SEM. Paired t-test was used to compare mean M6 responses to different light intensities (Figure 4).

**Figure 1.**
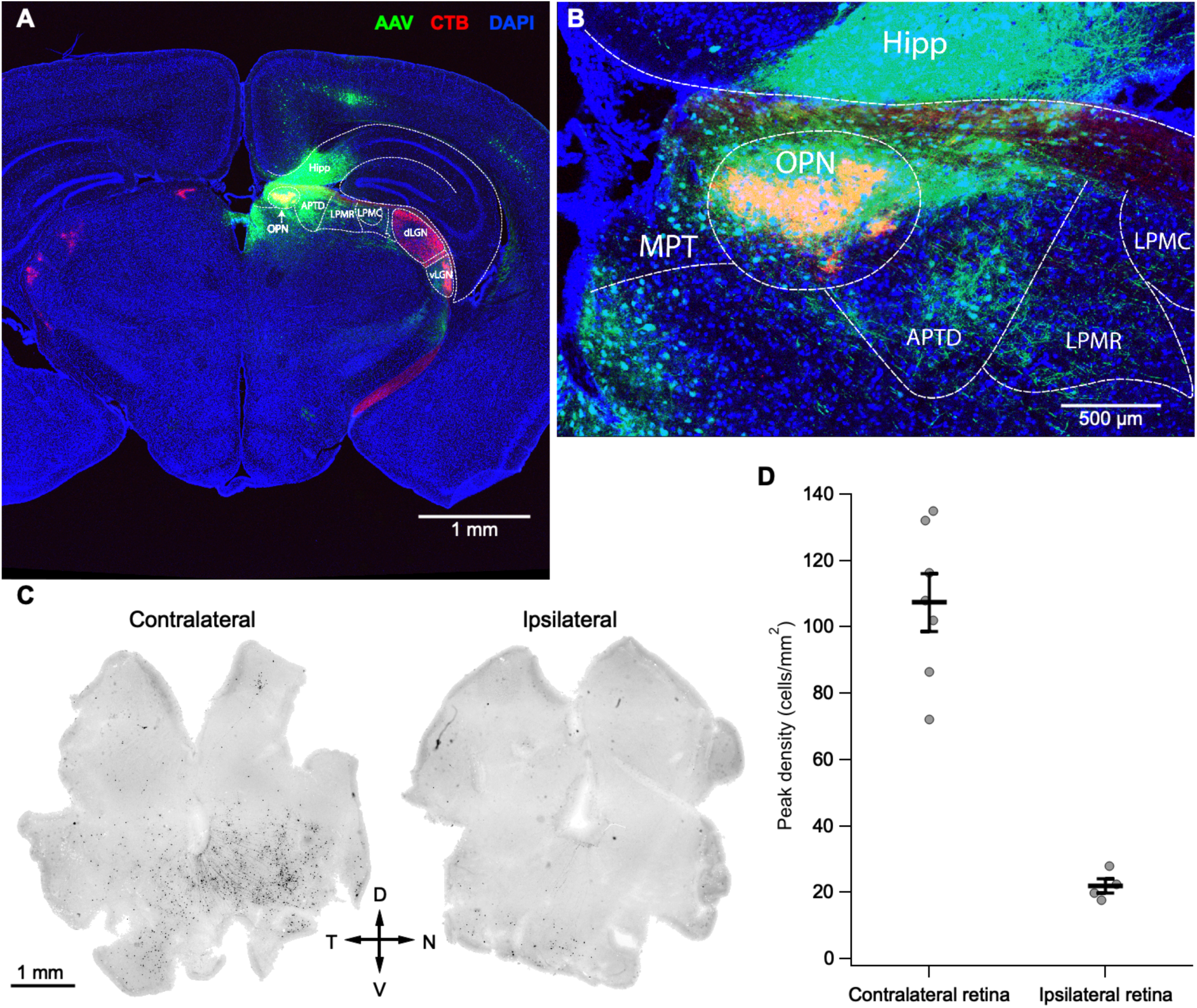
Retrograde labeling of OPN-projecting RGCs. **A**, Targeted injection of rostral aspect of OPN. Red: CTB-547 from contralateral intravitreal injection, demarcating retinorecipient regions. Green: AAVrg-CAG-GFP from injection, demarcating injection site. Green subcortical cell bodies away from injection site are from retrograde infection of the virus and indicate cells which project to OPN. Cortex and hippocampus along the injection tract are not retinorecipient (Morin and Studholme, 2014). Yellow: red from CTB-labeling of OPN, and green from virus injection, indicating target of the injection in the OPN. **B**, Higher magnification image of injection site in OPN. **C**, *Left*, Contralateral retina following 7 days of virus expression; *Right*, Ipsilateral retina. **D**, Peak density of labeled RGCs in contralateral and ipsilateral retina. Lines show mean and SEM of individual (grey) points from each retina. **Abbreviations:** APTD = anterior pretectal nucleus, dorsal aspect; dLGN = dorsal lateral geniculate nucleus; hipp = hippocampus; LPMC = lateral posterior thalamic nucleus, mediocaudal part; LPMR = lateral posterior thalamic nucleus, mediorostral part; MPT = medial pretectal nucleus; OPN = olivary pretectal nucleus; vLGN = ventral lateral geniculate nucleus (Paxinos and Franklin, 2012). D = dorsal; V = ventral; N = nasal; T = temporal.

**Figure 2.**
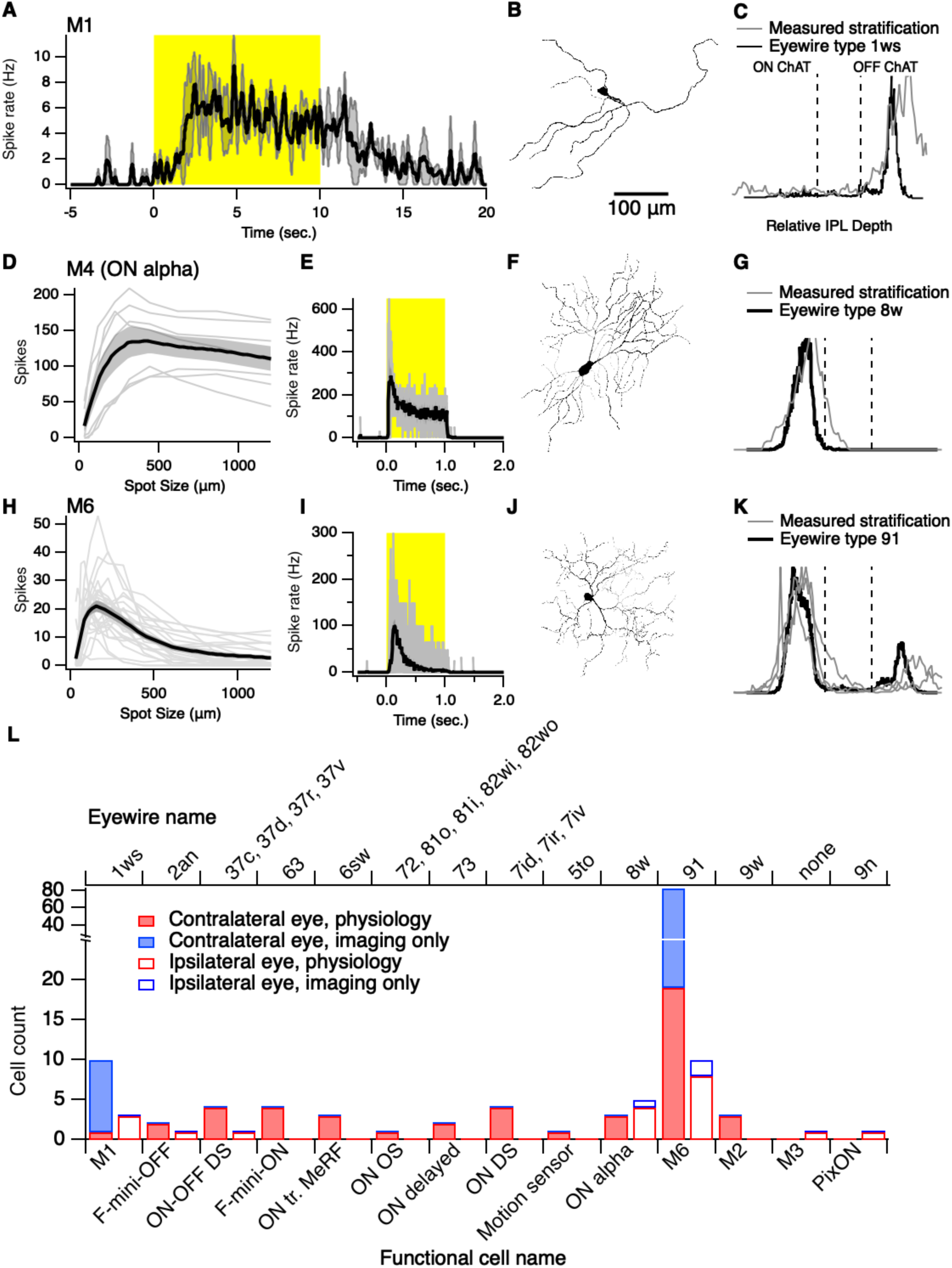
Typology of OPN-projecting RGCs. **A-C**, Physiology and morphology/stratification for M1 ipRGCs labeled by virus injection. **A**, Peri-stimulus time histogram of spiking response to presentation of full-field light step at 4744 r* (n = 2 M1 ipRGCs). Histograms were smoothed with a sliding weighted average. **B**, Inverted confocal image of one M1 ipRGC. **C**, Stratification of M1 ipRGC. ON and OFF refer to the depth of the ON and OFF ChAT bands. Gray, trace stratification of M1. Black, average stratification of type ‘1ws’, the putative M1 in the EyeWire data set, determined by electron microscopy (n = 2) (Bae et al., 2018). **D-G**, Physiology and morphology/stratification for ON alpha RGCs (M4 ipRGCs) labeled by virus injection. **D**, Spike count during stimulus presentation of spots of different diameters (average ± SEM). **E**, Peri-stimulus time histogram at preferred spot size of 313 μm. **F**, Inverted confocal image of one ON alpha RGC with physiology in D and E. **G**, Stratification as above. Gray, stratification of ON alpha RGC. Black, average of EyeWire type ‘8w’ (n = 4) **H-K**, As in **D-G**, but for M6 ipRGCs. **H**, n = 27 M6 ipRGCs. **I**, Preferred spot size = 224 μm, n = 27 M6 ipRGCs. **J**, Inverted confocal image of one M6 ipRGC with physiology in H and I. **K**, Gray, individual traces of M6 ipRGCs (n = 3). Black, average stratification of EyeWire type ‘91’ RGCs (n = 7). **L**, Bar graph of all identified RGCs from all retinas, identified on the basis of physiology or coarse morphology/stratification from epifluorescence imaging (contralateral retina, solid bars, n = 46 RGCs physiology, 73 cells imaging; ipsilateral retina, empty bars, n = 18 cells physiology, 3 cells imaging). **Abbreviations:** ON tr. MeRF = ON transient, medium receptive field (unpublished); OS = ON orientation selective; DS = direction selective.

**Figure 3.**
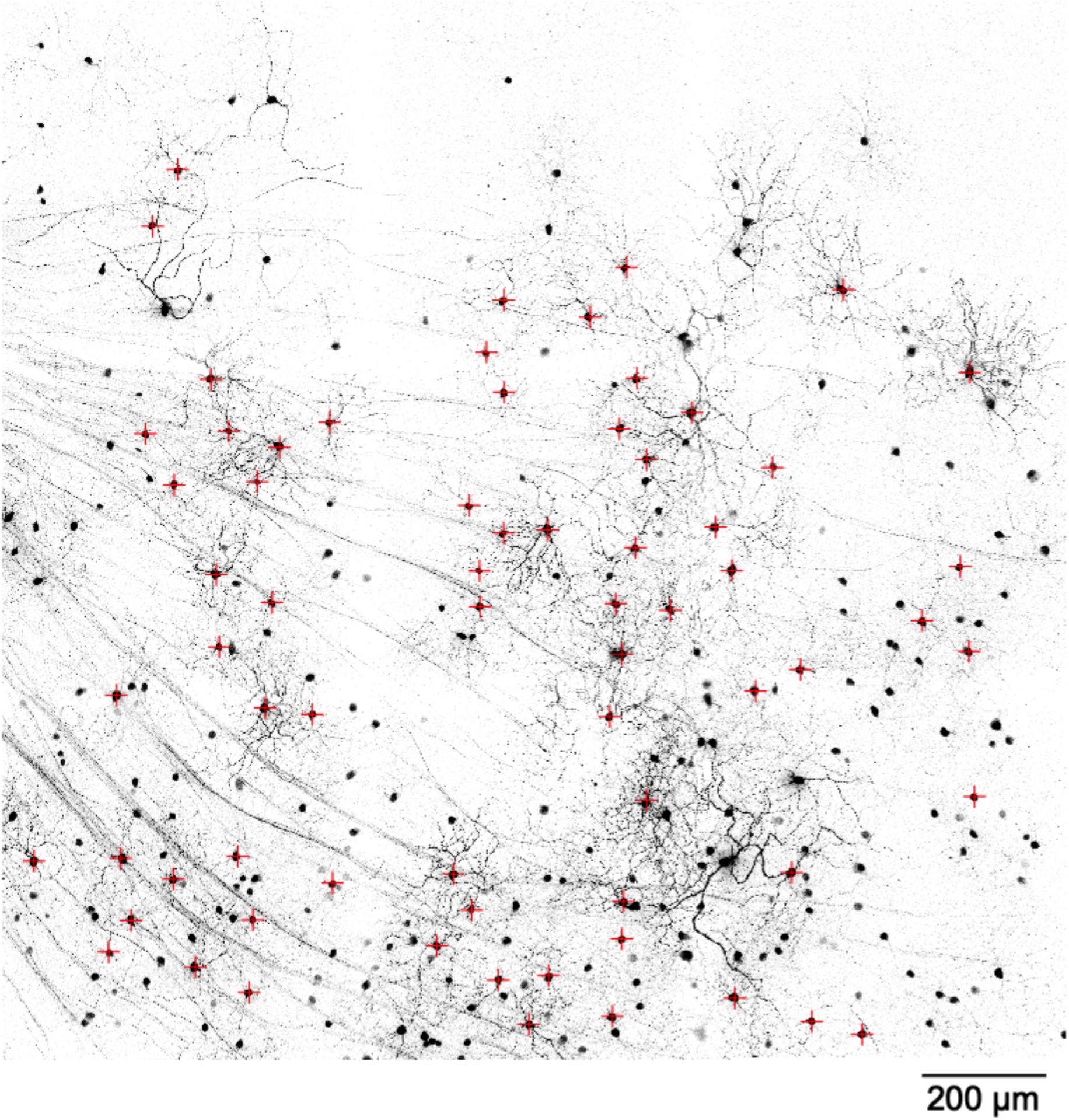
A dense population of M6 RGCs projects to the OPN. Image of the ventral-nasal region of a retina contralateral to the OPN injection. Red crosses indicate M6 RGCs confirmed by dendritic morphology in 3D.

**Figure 4.**
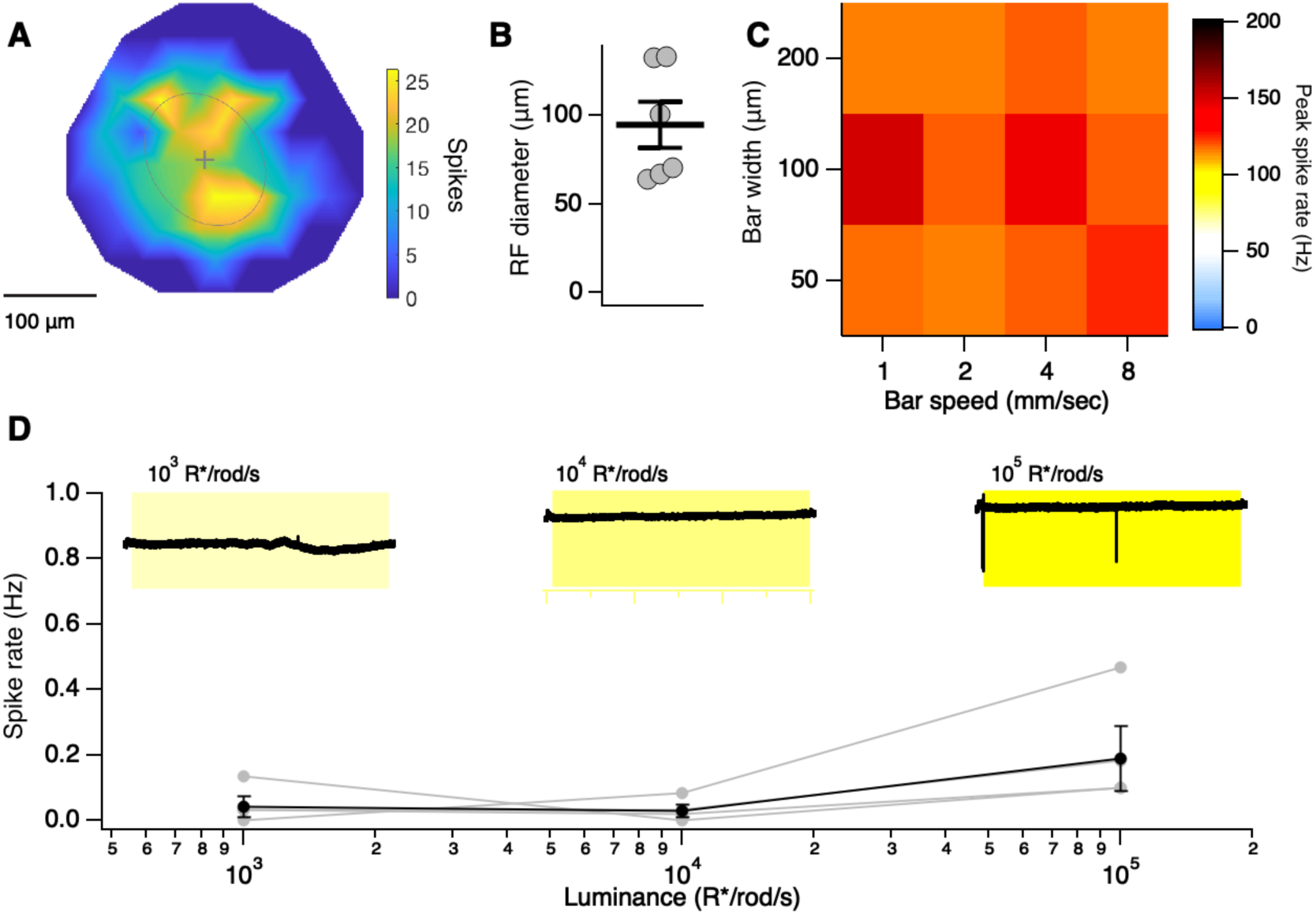
M6 RGCs detect small objects across a range of speeds but do not encode absolute luminance. **A**, Spiking receptive field of a representative M6 RGC. **B**, M6 RGC receptive field diameters. Lines indicate mean and SEM of individual cells (grey points). **C**, Peak spike rate of M6 RGCs to moving bars of various speeds and widths. Mean of 3 M6 RGCs. **D**, Mean spike rate of M6 RGCs over a 60 s light step. Data from individual cells (grey points) connected by lines. Black trace is the mean ± SEM. Insets show cell-attached traces from an example cell where brief periods of spiking were observed only at 10^5^ R*/rod/s. Yellow shading indicates the 60 s light stimulus. Recordings were all performed in the absence of synaptic blockers.

## Results

To identify the complement of RGC’s which project to the OPN, we made use of a virus engineered for retrograde transport which carried either the red fluorophore tdTomato or the green fluorophore GFP (rAAV2-CAG-tdTomato or rAAV2-CAG-GFP). Virus was injected stereotactically into the OPN, anterior to the nucleus of the optic tract (NOT) and the posterior pretectal nucleus (PPT), both nearby retinorecipient structures (Morin and Studholme, 2014). To verify the accuracy of the injection, we injected the retrograde tracer CTB into the eye contralateral to the brain injection site, thereby marking all retinorecipient brain regions with the far red dye Alexa647 (Figure 1A). The absence of CTB-labeled pretectal nuclei apart from the OPN near the brain injection site indicated that this structure was the only retinorecipient region in the area of the injection site (Figure 1A,B).

Injection of the OPN resulted in labeling of RGC somas and dendrites in the ventral-nasal portion of the contralateral retina, and sparse labeling in the ventral-temporal portion of the ipsilateral retina (Figure 1C), consistent with previous findings on the retinotopy of OPN (Scalia and Arango, 1979; Campbell and Lieberman, 1985; Young and Lund, 1998). We counted fluorescent somas within the densest 1 mm^2^ of labeled retina to measure the density of RGCs innervating OPN (Figure 1D). The average density in the contralateral eye was 107 ± 9 cells/mm^2^ (mean ± SEM, n = 7 retinas), and 22 ± 2 cells/mm^2^ in the ipsilateral eye (n = 4 retinas). These densities represent 3.25% and 0.66% of total RGC density, respectively (Jeon et al., 1998).

To determine the identity of the labeled RGCs, we performed targeted *ex vivo* electrophysiological recordings guided by two-photon imaging in both contralateral and ipsilateral retina (n = 7 contralateral retinas; n = 4 ipsilateral retinas, Figure 2). Cells were characterized primarily on the basis of their responses to spots of light of varying diameter, intensity and duration centered on their receptive fields (n = 65 cells, contralateral and ipsilateral combined). In some cases, typology was confirmed on the basis of dendritic morphology and stratification in the inner plexiform layer (IPL). Finally, the labeling of some cells from viral infection was of sufficient brightness and completeness that these cells could be identified by imaging on the basis of their morphology and coarse stratification without electrophysiology (n = 76 cells).

We found 14 types of RGCs that project to the OPN (Figure 2L). Intrinsically photosensitive retinal ganglion cells (ipRGCs) were the most abundant class of RGCs encountered, accounting for 82.9% (117/141) of all labeled RGCs. M1 ipRGCs have been shown previously to innervate the shell of the OPN, providing luminance information for the PLR (Chen et al., 2011). Consistent with these findings, we identified M1 ipRGCs (12/141), which responded to large spots of light with slow and sustained firing (Figure 2A), had a large, diffuse dendritic tree, and were stratified at the outer limit of the IPL (Figure 2B,C). We also identified a considerable proportion of M4 ipRGCs, also called ON-alpha RGCs, on the basis of their preference for spot sizes of large diameter, and their sustained firing and high spike rate, much higher than M1 RGCs (n = 8/141, Figure 2D,E). Morphologically, they had comparatively dense dendritic trees with large caliber primary dendrites, and they stratified in the ON layer (Figure 2F,G). The remaining identified types with sparse projections to OPN included both previously described types, like ON-OFF DS RGCs, and two unpublished types (ON tr. MeRF and Motion sensor) named in our lab’s large-scale classification project (Goetz et al., manuscript in preparation). The majority of these types were also ON RGCs, including ON direction-selective cells and F-mini-ON RGCs (7/9 non-ipRGC, Figure 2L).

By far the RGC type most commonly encountered in both contralateral and ipsilateral retinas was the recently identified M6 ipRGC, which comprised 66% of all identified RGCs (n = 93/141 cells, Figure 2L) (Quattrochi et al., 2019). Consistent with previous findings, M6 RGCs responded optimally to small spots of light and exhibited strong surround suppression to spot sizes beyond 200 μm (n = 27 cells, Figure 2H). Additionally, their light responses were transient, ending by 0.5 sec into the 1 s stimulus presentation (n =27, Figure 2I). They had small, moderately branched and spiny dendritic fields (Figure 2J), with bistratified, but primarily ON-stratified dendrites at the inner and outer margins of the IPL (n = 3 cells, Figure 2K), also consistent with the previous report (Quattrochi et al., 2019).

To estimate the proportion of the total retinal population of M6 RGCs labeled by our injections, we measured the density of labeled M6 somas in 1 mm^2^ of retina to be 23.3 cells/mm^2^. We compared this value to the density of the morphological match to the M6 RGC in the Eyewire museum, type ‘91’ (Bae et al., 2018), which also had a density of 23.3 cells/mm^2^. The close alignment of density values between the studies indicates that nearly all M6s were labeled by our injections, and thus, that the entire M6 population is likely to project to the OPN (Figure 3).

Because M6 RGCs account for such a large proportion of the retinal input to OPN, we performed additional functional experiments to explore what type of information they might transmit. Receptive field measurements with individual spots of light revealed a small receptive field (95 ± 13 μm^2^, n = 6 M6 RGCs, Figure 4A,B). Given that retinal ganglion cells vary in the range of speeds of motion to which they respond (Jacoby and Schwartz, 2017), we probed the speed sensitivity of M6 RGCs by recording their spike responses to moving bars of different speeds and widths. The peak firing rate was similar across all speeds tested, from 1 to 8 mm/sec and across widths from 50 to 200 μm (n = 3 M6 RGCs, Figure 4C).

Finally, given its classification as an ipRGC, we tested the ability of the M6 to encode absolute luminance by recording its spiking response to 60 s long, full-field steps of light of increasing intensity (Figure 4D). While previous work in synaptic blockers revealed a modest, melanopsin-driven current in the M6 in response to full-field light steps, because we were interested in the spiking information regarding absolute luminance the M6 might transmit to the OPN, these experiments were performed in the absence of synaptic blockers (Quattrochi et al., 2019). We found that this stimulus elicited only a weak spiking response even for a luminance of 10^5^ R*/rod/s, where most M1 ipRGCs show robust and sustained responses (Figure 4D, top traces) (Milner and Do, 2017; Lee et al., 2019). There was also no significant increase in spike rate across luminance in the M6 RGCs we measured (10^3^ vs. 10^5^ R*/rod/s, p = 0.24, Student’s paired t-test Fig. 4D, bottom). Therefore, M6 RGCs are unlikely to transmit absolute luminance information to the OPN, and any melanopsin-mediated current is likely only to have a modulatory effect on M6 function as in ON alpha (M4) ipRGCs (Sonoda et al., 2018).

## Discussion

Determining which of the more than 40 retinal ganglion cell types project to a given retinorecipient region is a vital step in elucidating how retinal output drives visually-guided behavior. We combined retrograde viral tracing with single-cell electrophysiology to identify the RGC types which project to the OPN. We identified 14 types of RGCs projecting to the OPN, nearly all of which were ON RGCs. Despite this variety of inputs, the majority were from a single type of RGC, the M6 RGC (Quattrochi et al., 2019). We confirmed that the M6 RGC has a transient ON response to light onset and responds optimally to small spot sizes of about 200 μm with nearly complete surround suppression for larger spots. We also showed that the M6 RGC responds robustly to the presence of moving objects across a wide range of speeds. Despite the presence of melanopsin, the M6 RGC does not modulate its maintained spiking output with changes in luminance, suggesting that any melanopsin-mediated current is likely modulatory in its effect on M6 physiology.

Given our result that much of the input to OPN conveys information about high spatial resolution rather than luminance, it is fair to speculate that this nucleus serves a function in addition to its well-established role in the pupillary light reflex. Consistent with this idea is the substantial connectivity from the OPN to other visual structures, including the superior colliculus, suprachiasmatic nucleus, intergeniculate leaflet, the ventral lateral geniculate nucleus, and the lateral posterior-pulvinar complex of the thalamus (Klooster et al., 1995; Moga and Moore, 1997). Interestingly, there is a retinal projection to the OPN in the blind mole rat in spite of the absence of an iris or ciliary body (Cooper et al., 1993). Other tracing studies have revealed that a subset of M1 ipRGCs connects with the shell of the OPN, and that the pupillary light reflex is severely attenuated when these cells are genetically ablated (Chen et al., 2011). Such work leaves open the possibility that other functions may be mediated by the core of the OPN, and projections to the aforementioned targets of OPN efferents.

Our findings suggest a number of experiments to illuminate M6 functionality and its role in visually-guided behavior mediated by the OPN. First, what is the function of melanopsin in the M6 if not to encode absolute luminance? Despite a small current, melanopsin activation leads to increased excitability in M4 (ON alpha) RGCs through a different transduction pathway than that in M1 RGCs (Sonoda et al., 2018). The nature of the melanopsin transduction pathway in M6 RGCs and its relationship to function remain open questions. Second, recordings from neurons within the OPN have centered around luminance sensitivity, revealing the presence of cells which increase spiking monotonically with increasing luminance (Clarke and Ikeda, 1985). Single-cell recording and imaging studies of the OPN with the presentation of moving objects could reveal additional visual sensitivity. Additionally, genetic silencing of retinal input to OPN could reveal behavioral effects beyond disruption of the PLR, perhaps even in a task involving small moving objects, like prey-capture (Hoy et al., 2016). Finally, It should be noted that our findings on M6 physiology are not inconsistent with a role in the PLR. Previous work has shown that rod photoreception is vital for the fast component of pupil constriction (Keenan et al., 2016). Therefore, It is possible that the M6 contributes to the fast kinetics of the PLR during its transient response to light onset. If an appropriately specific M6-only mouse line is created, silencing M6 RGCs while testing the PLR would test whether they play a role in this reflex.

## Acknowledgements

We acknowledge Tiffany Schmidt, J.C. Cang, and Yongling Zhu for their helpful comments on an earlier version of this manuscript.

## References

Baden T, Berens P, Franke K, Román Rosón M, Bethge M, Euler T (2016) The functional diversity of retinal ganglion cells in the mouse. Nature 529:345–350.

Bae JA, Mu S, Kim JS, Turner NL, Tartavull I, Kemnitz N, Jordan CS, Norton AD, Silversmith WM, Prentki R, Sorek M, David C, Jones DL, Bland D, Sterling ALR, Park J, Briggman KL, Seung HS, Eyewirers (2018) Digital Museum of Retinal Ganglion Cells with Dense Anatomy and Physiology. Cell 173:1293–1306.e19.

Campbell G, Lieberman AR (1985) The olivary pretectal nucleus: experimental anatomical studies in the rat. Philos Trans R Soc Lond B Biol Sci 310:573–609.

Chen S-K, Badea TC, Hattar S (2011) Photoentrainment and pupillary light reflex are mediated by distinct populations of ipRGCs. Nature 476:92–95.

Clarke RJ, Ikeda H (1985) Luminance and darkness detectors in the olivary and posterior pretectal nuclei and their relationship to the pupillary light reflex in the rat. I. Studies with steady luminance levels. Exp Brain Res 57:224–232.

Cooper HM, Herbin M, Nevo E (1993) Visual system of a naturally microphthalmic mammal: the blind mole rat, Spalax ehrenbergi. J Comp Neurol 328:313–350.

Hattar S, Kumar M, Park A, Tong P, Tung J, Yau K-W, Berson DM (2006) Central projections of melanopsin-expressing retinal ganglion cells in the mouse. J Comp Neurol 497:326–349.

Hoy JL, Yavorska I, Wehr M, Niell CM (2016) Vision Drives Accurate Approach Behavior during Prey Capture in Laboratory Mice. Curr Biol 26:3046–3052.

Jacoby J, Schwartz GW (2017) Three Small-Receptive-Field Ganglion Cells in the Mouse Retina Are Distinctly Tuned to Size, Speed, and Object Motion. J Neurosci 37:610–625.

Jeon C-J, Strettoi E, Masland RH (1998) The Major Cell Populations of the Mouse Retina. The Journal of Neuroscience 18:8936–8946 Available at: http://dx.doi.org/10.1523/jneurosci.18-21-08936.1998.

Keenan WT, Rupp AC, Ross RA, Somasundaram P, Hiriyanna S, Wu Z, Badea TC, Robinson PR, Lowell BB, Hattar SS (2016) A visual circuit uses complementary mechanisms to support transient and sustained pupil constriction. Elife 5 Available at: http://dx.doi.org/10.7554/eLife.15392.

Klooster J, Beckers HJM, G F J, van der Want JJL (1993) The peripheral and central projections of the Edinger-Westphal nucleus in the rat. A light and electron microscopic tracing study. Brain Research 632:260–273 Available at: http://dx.doi.org/10.1016/0006-8993(93)91161-k.

Klooster J, G F J, Müller LJ, van der Want JJL (1995) Efferent projections of the olivary pretectal nucleus in the albino rat subserving the pupillary light reflex and related reflexes a light microscopic tracing study. Brain Research 688:34–46 Available at: http://dx.doi.org/10.1016/0006-8993(95)00497-e.

Lee SK, Sonoda T, Schmidt TM (2019) M1 Intrinsically Photosensitive Retinal Ganglion Cells Integrate Rod and Melanopsin Inputs to Signal in Low Light. Cell Rep 29:3349–3355.e2.

Martersteck EM, Hirokawa KE, Evarts M, Bernard A, Duan X, Li Y, Ng L, Oh SW, Ouellette B, Royall JJ, Stoecklin M, Wang Q, Zeng H, Sanes JR, Harris JA (2017) Diverse Central Projection Patterns of Retinal Ganglion Cells. Cell Rep 18:2058–2072.

Milner ES, Do MTH (2017) A Population Representation of Absolute Light Intensity in the Mammalian Retina. Cell:1–12.

Moga MM, Moore RY (1997) Organization of neural inputs to the suprachiasmatic nucleus in the rat. J Comp Neurol 389:508–534.

Morin LP, Studholme KM (2014) Retinofugal projections in the mouse. J Comp Neurol 522:3733–3753.

Paxinos G, Franklin KBJ (2012) Paxinos and Franklin’s the Mouse Brain in Stereotaxic Coordinates. Academic Press.

Quattrochi LE, Stabio ME, Kim I, Ilardi MC, Michelle Fogerson P, Leyrer ML, Berson DM (2019) The M6 cell: A small-field bistratified photosensitive retinal ganglion cell. J Comp Neurol 527:297–311.

Rousso DL, Qiao M, Kagan RD, Yamagata M, Palmiter RD, Sanes JR (2016) Two Pairs of ON and OFF Retinal Ganglion Cells Are Defined by Intersectional Patterns of Transcription Factor Expression. Cell Reports 15:1930–1944 Available at: http://dx.doi.org/10.1016/j.celrep.2016.04.069.

Scalia F, Arango V (1979) Topographic organization of the projections of the retina to the pretectal region in the rat. J Comp Neurol 186:271–292.

Sonoda T, Lee SK, Birnbaumer L, Schmidt TM (2018) Melanopsin Phototransduction Is Repurposed by ipRGC Subtypes to Shape the Function of Distinct Visual Circuits. Neuron 99:754–767.e4.

Sümbül U, Song S, McCulloch K, Becker M, Lin B, Sanes JR, Masland RH, Sebastian Seung H (2014) A genetic and computational approach to structurally classify neuronal types. Nature Communications 5 Available at: http://dx.doi.org/10.1038/ncomms4512.

Trejo LJ, Cicerone CM (1984) Cells in the pretectal olivary nucleus are in the pathway for the direct light reflex of the pupil in the rat. Brain Res 300:49–62.

Young MJ, Lund RD (1998) The retinal ganglion cells that drive the pupilloconstrictor response in rats. Brain Res 787:191–202.

Campbell, G., and A. R. Lieberman. 1985. “The Olivary Pretectal Nucleus: Experimental Anatomical Studies in the Rat.” Philosophical Transactions of the Royal Society of London. Series B, Biological Sciences 310 (1147): 573–609.

Martersteck, Emily M., Karla E. Hirokawa, Mariah Evarts, Amy Bernard, Xin Duan, Yang Li, Lydia Ng, et al. 2017. “Diverse Central Projection Patterns of Retinal Ganglion Cells.” Cell Reports 18 (8): 2058–72.

Morin, Lawrence P., and Keith M. Studholme. 2014. “Retinofugal Projections in the Mouse.” The Journal of Comparative Neurology 522 (16): 3733–53.

Quattrochi, Lauren E., Maureen E. Stabio, Inkyu Kim, Marissa C. Ilardi, P. Michelle Fogerson, Megan L. Leyrer, and David M. Berson. 2019. “The M6 Cell: A Small-Field Bistratified Photosensitive Retinal Ganglion Cell.” The Journal of Comparative Neurology 527 (1): 297–311.

Rousso, David L., Mu Qiao, Ruth D. Kagan, Masahito Yamagata, Richard D. Palmiter, and Joshua R. Sanes. 2016. “Two Pairs of ON and OFF Retinal Ganglion Cells Are Defined by Intersectional Patterns of Transcription Factor Expression.” Cell Reports. https://doi.org/10.1016/j.celrep.2016.04.069.

Scalia, F., and V. Arango. 1979. “Topographic Organization of the Projections of the Retina to the Pretectal Region in the Rat.” The Journal of Comparative Neurology 186 (2): 271–92.

